# Conjugative dissemination of plasmids in rapid sand filters: a trojan horse strategy to enhance pesticide degradation in groundwater treatment

**DOI:** 10.1101/2020.03.06.980565

**Authors:** Rafael Pinilla-Redondo, Asmus Kalckar Olesen, Jakob Russel, Lisbeth Elvira de Vries, Lisbeth Damkjær Christensen, Sanin Musovic, Joseph Nesme, Søren Johannes Sørensen

## Abstract

The supply of clean water for human consumption is being challenged by the appearance of pesticide pollutants in groundwater ecosystems. Biological rapid sand filtration is a commonly employed method for the removal of organic and inorganic impurities in water which relies on the degradative properties of microorganisms for the removal of diverse contaminants, including pesticides. Although sustainable and relatively inexpensive, the bioremediation capabilities of rapid sand filters vary greatly across waterworks. Bioaugmentation efforts with degradation-proficient bacteria have proven difficult due to the inability of the exogenous microbes to stably colonize the sand filters. Pesticide degrading genes, however, are often encoded naturally by plasmids—extrachromosomal DNA elements that can transfer between bacteria—yet their ability to spread within rapid sand filters have remained unknown. To evaluate the potential use of plasmids for the dissemination of pesticide degrading genes, we examined the permissiveness of rapid sand filter communities towards four environmental transmissible plasmids; RP4, RSF1010, pKJK5 and TOL (pWWO), using a dual-fluorescent bioreporter platform combined with FACS and 16S rRNA gene amplicon sequencing. Our results reveal that plasmids can transfer at high frequencies and across distantly related taxa from rapid sand filter communities, emphasizing their suitability for introducing pesticide degrading determinants in the microbiomes of underperforming water purification plants.

## Introduction

Safe drinking water is becoming a scarce commodity in many parts of the world and several countries rely on groundwater systems for its supply (Navarrete *et al.*, 2008). However, with the advent of modern agriculture, urbanization, and other anthropogenic practices, groundwater reservoirs are threatened by the leaching of chemical pollutants and their toxic degradation products (Fenner *et al.*, 2013). In particular, pesticide contamination has been reported to be widespread and recalcitrant among subsoil aquifers, thus constituting an increasing environmental and human health concern (Malaguerra *et al.*, 2012; Pérez-Lucas *et al.*, 2019).

Biological rapid sand filters (sand filters) are commonly employed for the treatment of raw groundwater. Apart from efficiently removing large suspended particles and other impurities, sand filters are involved in the biodegradation of organic matter and ammonium removal —processes that heavily rely on the resident bacterial communities (Lee *et al.*, 2014). Importantly, this water purification approach constitutes a relatively cost-effective and environmentally friendly practice in contrast to other more advanced technologies, including reverse osmosis (Lee, Howe and Thomson, 2012), advanced oxidation (Suty, de Traversay and Cost, 2004), or granular activated carbon (Snyder *et al.*, 2007). The microbial communities in sand filters are not naturally adapted for the removal of pesticides, leading most common pesticide residues in the groundwater (e.g. BAM, Bentazone, Atrazine etc.) to filter though into the drink water. To mitigate this problem, the amendment of sand filters with bacteria harboring the desired catabolic genes has been proposed as a means to enhance the degrading potential of underperforming waterworks (Benner *et al.*, 2013). Nonetheless, such bioaugmentation strategies have met little success because of the low retention times of the introduced strains, a problem that is mainly attributed to the colonization resistance (biological barrier effect) exerted by the indigenous sand filter microbial communities (Benner *et al.*, 2013; Albers *et al.*, 2015).

While the genes involved in the degradation of pesticides can sometimes be encoded on the chromosomes of bacteria, they are often found to be carried naturally by mobile genetic elements (MGEs) (Springael and Top, 2004; Dunon *et al.*, 2013; Dealtry *et al.*, 2014). Among these, conjugative and mobilizable plasmids are of particular interest since they are widely recognized as effective vectors for the dissemination of genetic traits across microbiomes (Norman, Hansen and Sørensen, 2009). Indeed, certain plasmids have been reported to exhibit remarkably promiscuous properties, readily transferring into a wide range of phylogenetically distant taxa (Jain and Srivastava, 2013; Klümper *et al.*, 2015). Consequently, the delivery of plasmids harboring pesticide degrading genes presents itself as a promising alternative to strain-based bioaugmentation strategies. Since the establishment of a plasmid donor strain within a microbial community is not a prerequisite for transfer (Licht and Wilcks, 2005), this approach could serve as a “trojan horse strategy” to enhance the pesticide degrading capabilities of sand filters, while bypassing the aforementioned colonization resistant hurdle. In order to assess the potential of plasmid-derived bioremediation approaches, a better understanding of the permissiveness of sand filter bacterial communities to the dissemination of exogenous plasmids is needed, yet to date, the transfer frequencies and host ranges of conjugative plasmids within sand filter microbiomes have remained unexplored.

Here, we investigate the potential permissiveness of bacterial communities originating from sand filters of water purification plants from three different geographic locations in Denmark (Kerteminde, Herning, and Bregnerød) to the transfer of four fluorescently-tagged environmental plasmids (pKJK5, TOL, RSF1010, and RP4). To address this question, we challenged the recipient sand filter communities with each plasmid using *Pseudomonas putida* as the plasmid-donor strain. We define community permissiveness here as the ability for native bacteria in the recipient rapid sand filter community to receive and express a reporter gene harbored by our plasmids. Plasmid transfer was monitored utilizing a well-established dual-fluorescent bioreporter platform in combination with high-throughput fluorescence activated cell sorting (FACS) and 16S rRNA gene amplicon sequencing of transconjugant and recipient cells (Pinilla-Redondo *et al.*, 2018). We report the first estimates of plasmid dissemination frequencies and host ranges within sand filter microbiomes, revealing high plasmid transfer frequencies and broad dissemination across bacterial taxa. Taken together, our data demonstrate the potential biotechnological application of natural plasmids for delivering pesticide-removal capabilities to microbial sand filter communities.

## Methods

### Strains, plasmids and sand filter recipient communities

The bacterial strains and plasmids and their relevant characteristics are listed in Table 1. Pseudomonas putida KT2440 (chromosomally tagged by *lacI_q_-Plpp-mCherry*), carrying either pKJK5, RP4, TOL or RSF1010, was used as the plasmid donor strain in the mating experiments. Sand filter sediments were sampled from water purification plants in Denmark: Kerteminde (55°25’47.5”N 10°38’18.2”E), Herning (56°08’48.5”N 8°56’33.7”E) and Bregnerød (55°48’51.9”N 12°22’36.2”E), representing different geographically distinct rapid sand filter microbial communities (Supplementary Fig. S1). The indigenous sand filter bacteria (T0) were extracted using the Nycodenz-gradient extraction method (Musovic *et al.*, 2010). Briefly, the sand material was grinded with a mortar in 50 mM TTSP (tetrasodium pyrophosphate buffer) and layered on top of the Nycodenz solution (Nycomed Pharma, Norway; 1.3 g/ml) prior to centrifugation (8,500 g for 15 min). The upper and intermediate phases containing the bacterial cells were collected, resuspended in x5 volumes of PBS and stored at 5 C until use. The donor strains were routinely grown in LB broth (10 g tryptone, 5 g yeast extract, and 4 g NaCl), using the appropriate antibiotics added at final concentrations of tetracycline (20 μg/ml), kanamycin (50 μg/ml). The recipient communities were grown in 5% TSB overnight at 30 °C and 150 rpm to enrich for the culturable fraction of bacteria in this environment prior to mating.

**Table 1.**
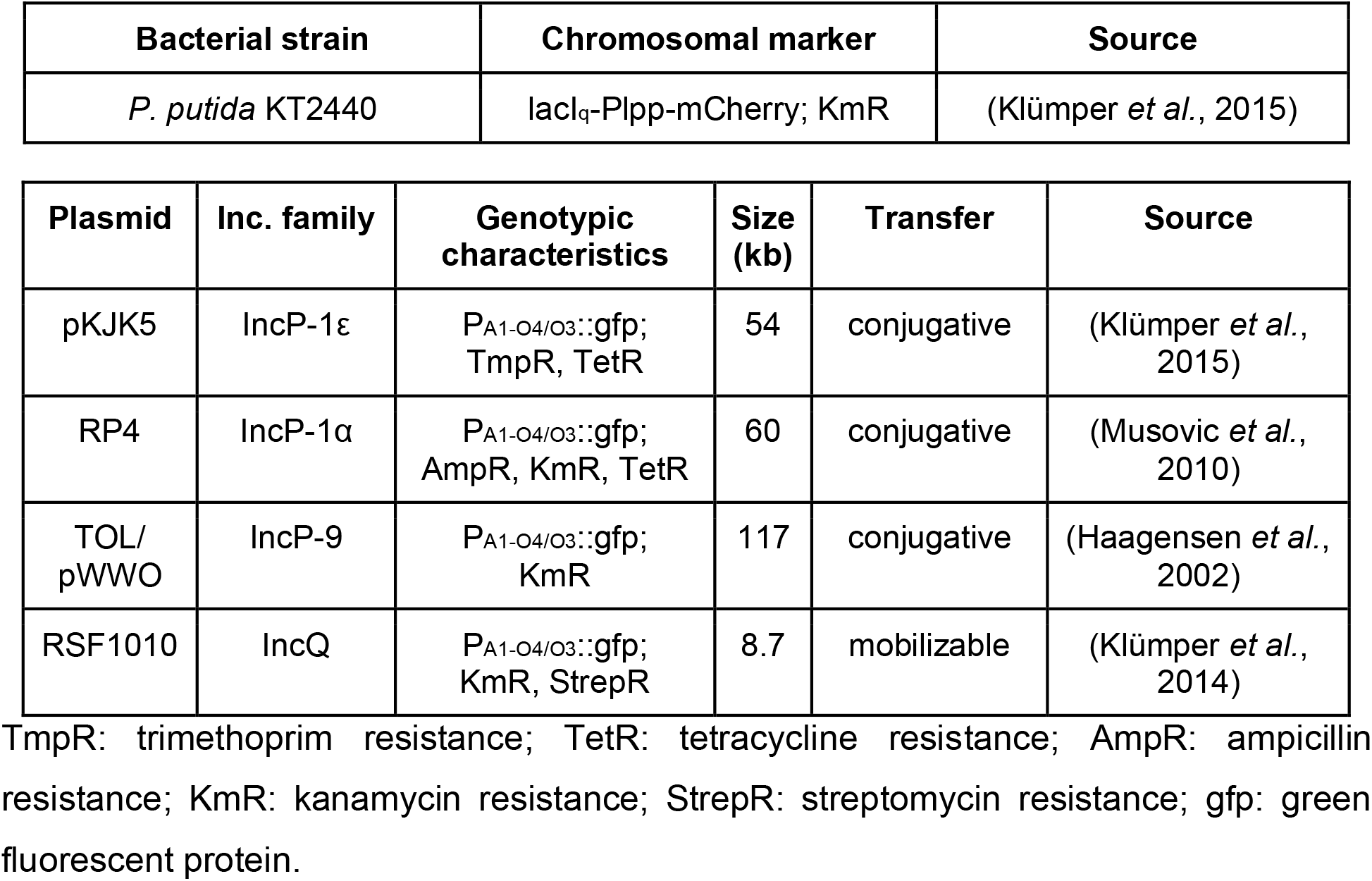

### Solid-surface plasmid conjugation assay

The permissiveness of the sand filter recipient communities towards the exogenous conjugative/mobilizable plasmids was tested using a modified version of a solid-surface meta-parental mating setup described previously (Klümper *et al.*, 2015). Following this approach, plasmids are tracked through a inserted *gfp* marker controlled by a *lacIq* repressible promoter. The donor strain additionally harbors a chromosomal *lacIq-Plpp-mcherry* insertion. Thus, in plasmid donor cells constitutive LacI production results in repression of the plasmid-encoded GFP, while the constitutive mCherry expression renders the cells red. The *gfp*-tagged plasmids, however, upon transfer into natural sand filter recipients are able to express GFP because these bacteria lack the *lacIq* insert found in the donor, thus ensuring a green fluorescent phenotype for transconjugant cells (Pinilla-Redondo *et al.*, 2018).

Filter matings were carried out by challenging the extracted sand filter recipient communities with the four plasmid-donor strain combinations independently, in triplicates, as well as performing negative control matings with the recipient communities and donor strains grown alone on the filters. Donor and recipient cell suspensions were adjusted to OD600 = 0.5 and 100 μl of each mixed at a 1:1 ratio. The resulting suspension was transferred onto sterile 0,2 μm nitrocellulose filters (Advantec) that were placed over 10% TSB agar media (Sigma Aldrich) without antibiotic selection. The area of the filter exposed was estimated to be 54 mm2, resulting in an initial cell count of approximately 3.6×10^5 cells/mm2. When dry, plates were incubated at 30°C for 24 h and filters were washed with 5 ml PBS to recover cells for FACS analysis (cell counting and sorting). Mating samples were kept at 4°C after recovery from filters and analyzed within a period of 3-4 days. The transfer efficiencies were calculated as the number transconjugant cells (T) divided by the geometric mean of the number of donor and recipient cells (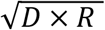) after mating (Sørensen and Jensen, 1998).

### FACS analysis

Flow cytometric detection of cells was performed by using a FACSAria Illu (Becton Dickson Biosciences, San Jose, CA, USA). The following technical settings were employed: A 70 μm nozzle and sheath fluid pressure of 70 psi; GFP was excited by a 488 nm laser (20mW) and detected on the FITC-A channel with a bandpass filter of 530/30 nm; mCherry was excited with a 561 nm laser (50mW) and detected on the PE-Texas Red-A channel with a bandpass filter of 610/20 nm. Detection thresholds were set to 200 for FSC and SSC. BD FACSDiva software v.6.1.3 was used for operating and analyzing the results.

Bivariate contour plots of particle FSC vs. SSC areas were employed to build a gate around the total bacterial population, excluding the background noise. Green and red fluorescent bacterial cells were gated on bivariate contour plots using the area of FITC vs. the area of PE-Texas red. The detection gates used in this study are depicted in Supplementary Figure S5. Donor, recipient and transconjugant counts were made with the “mCherry” (red), “green-non-red” and “non-red” gates, respectively. Flow cytometric analysis was performed by diluting filter mating samples in PBS to a cell count of 1000-3000 threshold evt/s, processed at flow rate 1. A total of 100,000 bacterial events were recorded for each mating outcome. Sorting of cells was performed into 5 mL sterile polypropylene round-bottom tubes (Falcon by Corning, USA) containing 0.5 mL of PBS. Because transconjugant cells often comprise less than 1% of the total cell population, we performed a preliminary sorting round as an enrichment step for transconjugant cells, as described in (Klümper *et al.*, 2015). First, 200,000-500,000 target transconjugant events were sorted using a flow rate of ~15,000 events/sec and employing the “yield/recovery” settings (both the interrogated drop and the drop adjacent to the target particle are sorted). Subsequently, a second more restrictive sorting step employing “purity” settings and a threshold rate <3000 events/s, was carried out to sort high purity transconjugant cells (any target events falling close to any non-target events are not sorted). In the second sorting round 20,000 cells were isolated from all filter-mating combinations. Sorted cells were then prepared for subsequent deep amplicon sequencing of 16S rRNA genes.

### Nucleic acids extraction

Microbial community profiling was carried out by 16S rRNA gene amplicon sequencing for the original sand filter community, hereafter referred to as “T0” (Supplementary Fig. S3). DNA was extracted using the Nucleospin soil kit (Macherey-Nagel) following the manufacturer’s instructions, using the lysis buffer SL1 and a bead-beating mechanical lysis step performed on a FastPrep-24 (MP Biomedicals) tissue homogenizer at 6 m/s for 30 sec. After filter mating, sorted transconjugant and recipient cells (referred to as “filter”) were pelleted by centrifugation at 10,000 x g for 30 min and cell lysis and DNA extraction were carried out in the thermal cycler, following the protocol detailed by the GenePurgeDirect (Nimagen) direct PCR kit.

### 16S rRNA gene amplicon sequencing

Sequencing libraries were prepared using a dual-PCR setup as described previously (Nunes *et al.*, 2016), targeting variable regions V3 and V4 of the 16S rRNA gene, approximately 460 bp. In the first step primers Uni341F (5’-CCTAYGGGRBGCASCAG-3’) and Uni806R (5’-GGACTACNNGGGTATCTAAT-3’) originally published by (Yu *et al.,* 2005) and modified as described in (Sundberg *et al.*, 2013) were used. In a second PCR reaction step the primers additionally included Illumina specific sequencing adapters and a unique combination of index for each sample (Nunes et al., 2016). PCR reactions were performed in 25 μl reactions using PCRBIO HiFi polymerase and 2 μL template DNA, following manufacturers’ instructions and the following program: 95°C for 1 min followed by 30 or 15 cycles (for, respectively, PCR1 or PCR2) of 95°C for 15 secs; 56°C for 15 secs and 72°C for 30 secs. After both PCR reactions, amplicon products were purified using HighPrep™ PCR Clean Up System (AC-60500, MagBio Genomics Inc., USA) using a 0.65:1 (beads:PCR reaction) volumetric ratio to remove DNA fragments below 100 bp in size. Samples were normalized using SequalPrep Normalization Plate (96) Kit (Invitrogen, Maryland, MD, USA) and pooled using 5 μl volume of each. The final pool volume was reduced to concentrate the sequencing library using the DNA Clean and Concentrator™-5 kit (Zymo Research, Irvine, CA, USA). The pooled library concentration was determined using the Quant-iT™ High-Sensitivity DNA Assay Kit (Life Technologies) following the specifications of the manufacturer. Before library denaturation and sequencing, the final pool concentration is adjusted to 4 nM before library denaturation and loading. Amplicon sequencing was performed on an Illumina MiSeq platform using Reagent Kit v2 [2×250 cycles] (Illumina Inc., CA, US). The MiSeq Controller Software Casava 1.8 (Illumina, USA) was used for sequence demultiplexing and the pair-end FASTQ output files were used for the downstream sequencing analysis. Raw sequence reads were first trimmed of primer sequences used in first PCR using cutadapt (Martin, 2011) and only read pairs for which both primers were found are retained for subsequent analysis. Primer trimmed sequences are then merged, clustered in OTU using UPARSE-OTU algorithm (Edgar, 2013) using a 97% pairwise sequence similarity threshold. The taxonomic annotation of each cluster representative sequence was performed using mothur (Schloss *et al.*, 2009) using the Ribosomal Database Project database trainset 16 (Cole *et al.*, 2014) (https://www.mothur.org/wiki/RDP_reference_files). An approximate maximum likelihood phylogenetic tree was built with FastTree (Price, Dehal and Arkin, 2010), based on alignment of all reference OTU cluster sequences obtained with mothur *align.seqs*.

### Sequence and data analyses

General handling of the data was carried out in R (R Development Core Team 3.0.1., 2019), through the following R packages: phyloseq (McMurdie and Holmes, 2013), reshape2 (Wickham, 2007), stringr (Wickham, 2019), dplyr (Wickham *et al.*, 2019) and plyr (Wickham, 2011). The prevalence method (threshold = 0.25) of the decontam package (Davis *et al.*, 2018) was used to remove potential contaminants from the dataset, removing 3.29 % of the total reads. The COEF package (Russel, 2019a) was used to remove OTUs that were not present in at least 2 out of 3 sample replicates across the whole dataset. Additionally, OTU reference sequences with a length below 380 were removed. Rarefied data, normalized to 15,000 reads per sample, were used for analysis of presence/absence of OTUs across samples. Non-rarefied data were used for analysis of relative abundances unless otherwise stated. Moreover, OTUs that were not present in a minimum of 2 out of 3 replicates per transconjugant sample group were removed. The “T0” samples describing the original sand filter community from Herning were removed from analyses due to indications of a technical error. The ggplot2 (Wilkinson, 2011) and ggpupr (Kassambara, 2018) packages were used for data visualization, and colors were adjusted using RColorBrewer (Neuwirth, 2014). Faith’s phylogenetic diversity metric (Faith, 1992) was calculated with the PhyloMeasures package (Tsirogiannis and Sandel, 2016) via the metagMisc package (https://github.com/vmikk/metagMisc). The statistical software package R (R Development Core Team 3.0.1., 2019) was used for analysis of variance (ANOVA) and to calculate Tukey honest significant differences. Weighted Unifrac distances (Lozupone and Knight, 2005) were calculated and plotted with the phyloseq package (McMurdie and Holmes, 2013). Permutational multivariate analysis of variance (PERMANOVA) tests were done with the vegan package (Oksanen *et al.*, 2019); using 999 permutations. Venn diagrams were constructed with the eulerr package (Larson *et al.*, 2018), through the MicEco package (Russel, 2019b). Heatmap plotting was made with the pheatmap package (Kolde, 2015) and differential abundance testing analyses were made with the DAtest package (Kolde, 2015). The taxonomic composition of transconjugant pools across plasmid-donor and sand filter recipient community combinations (Figure 2) were made using the iTOL webtool (Letunic and Bork, 2019), and the phylogenetic tree used as input was written from the phyloseq object using the ape package (Paradis and Schliep, 2019).

### Data availability

All raw sequence reads data have been deposited on EBI-SRA under BioProject accession number PRJEB36794.

## Results and Discussion

### Transfer efficiencies vary across sand filter recipient communities and plasmid-donor combinations

As a first step to evaluate the feasibility of plasmids for the introduction of pesticide degrading genes into sand filters, we explored the potential uptake of exogenous plasmids by bacterial communities originating from this environment. For this purpose, we conducted meta-parental matings in which bacteria extracted from 3 different waterworks in Denmark (Kerteminde, Herning, Bregnerød; Supplementary Figure S1) were challenged with a donor strain carrying one of the following GFP-tagged plasmids: pKJK5, RP4, RSF1010, TOL. In order to best assess the possibility of transfer, special attention was given to the choice of a relevant plasmid donor strain and plasmids. *Pseudomonas putida* was selected as the plasmid-donor model organism because it is a common dweller of water and soil environments (Chen *et al.*, 2014). Four different plasmids were chosen on the basis of 1) their intrinsic horizontal transferability properties—by being either conjugative (self-transmissible) or mobilizable (non-self-transmissible, yet able to use the conjugative machinery of a co-occurring conjugative plasmid for transfer) and, 2) their ability to replicate in hosts that naturally feed into drinking water reservoirs (eg. *Pseudomonas*) (Table 1).

The mean transfer efficiencies observed for the tested plasmids ranged from 10^-4^ to 10^-1^ across sand filter communities and plasmid-donor combinations (Figure 1), indicating that bacteria originating from these environments are permissive to the uptake of exogenous plasmids. It is worth considering that the efficiency of transfer events in nature may be lower than the levels detected here due to the potential bias associated with the clonal expansion of early transconjugant cells and because our solid-surface mating conditions are designed to maximize bacterial metabolic activity and cell-to-cell contact between donor and recipient cells. All things considered, these results indicate that the tested plasmids effectively permeate in the native recipient communities at the tested conditions of transfer.

**Figure 1.**
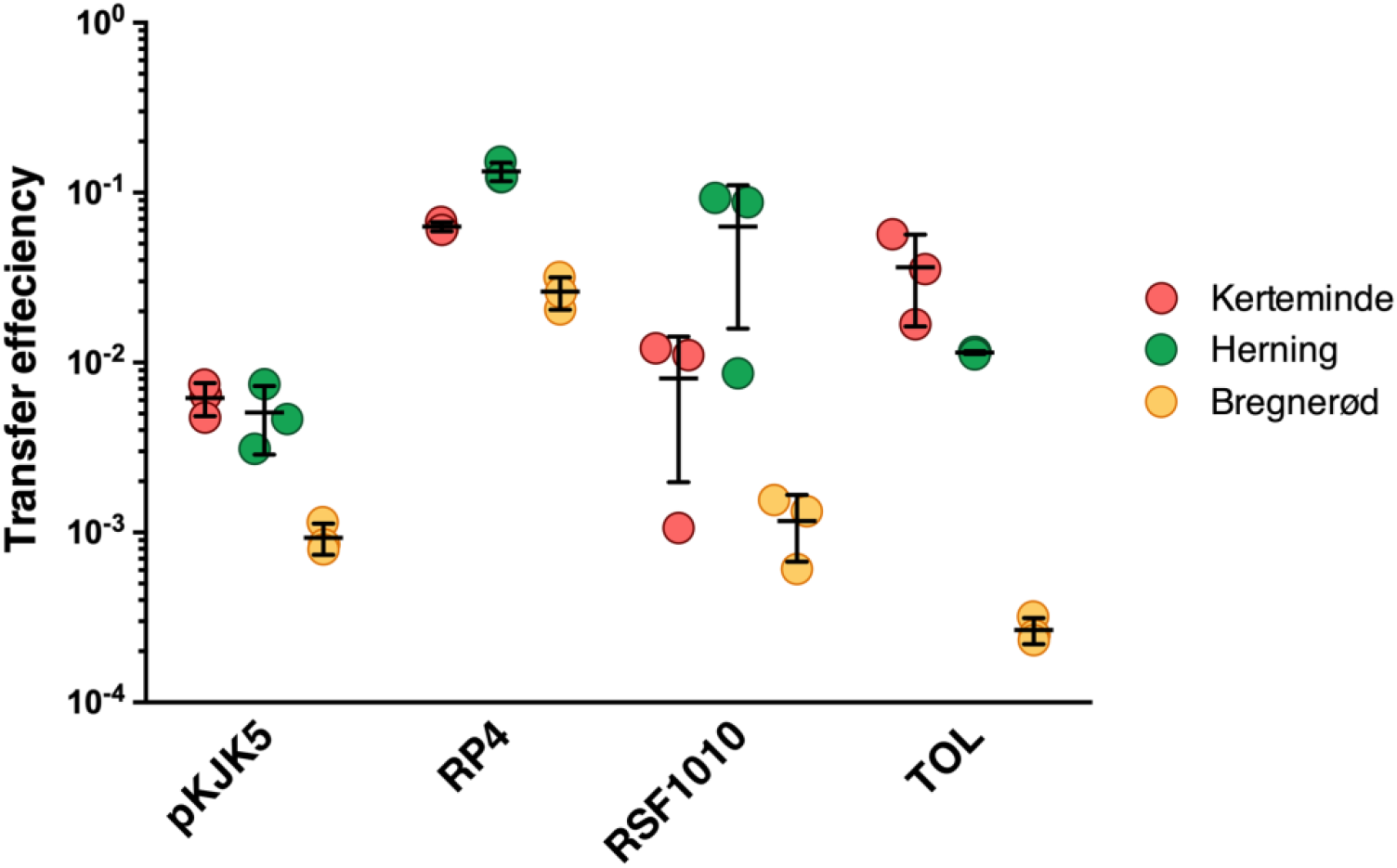
The efficiency of plasmid transfer varies across rapid sand filter communities and plasmid-donor combinations. Transfer efficiencies of pKJK5, RP4, RSF1010 and TOL resulting from filter matings using *P. putida* as the plasmid donor and recipient communities originating from sand filter waterworks in Kerteminde, Herning, and Bregnerød. Transfer efficiencies are expressed as the number of transconjugants divided by the geometric mean of donor and recipient cells, for each mating outcome. Error bars indicate the standard deviation of the mean for three independent filter mating replicates (Supplementary Table 1).

Overall, the four plasmids showed marked differences in their transfer efficiencies between sand filters, implying the existence of specific plasmid transfer bottlenecks across recipient communities. Interestingly, Bregnerød exhibited lower transfer efficiencies in comparison to Herning and Kerteminde for all tested plasmid-donor combinations, and RP4 showed more consistent high transfer efficiencies across sand filter communities (Figure 1). These results may reflect differences in the availability of suitable donor-recipient encounters and/or nuances in compatibility between plasmids and the genomic backgrounds they sample across sand filter bacterial communities. A plasmid’s entry and stable maintenance within a new host can be influenced by the host’s innate and adaptive barriers against incoming foreign DNA (Thomas and Nielsen, 2005; Bernheim and Sorek, 2019) and conflicts with co-resident MGEs. While the former includes bacterial defense systems such restriction-modification (Vasu and Nagaraja, 2013), CRISPR-Cas (Barrangou and van der Oost, 2012), and Wadjet (Doron *et al.,* 2018), the latter involves plasmid incompatibility issues with indigenous plasmids (Thomas, 2018) or may result from plasmid-plasmid competition dynamics, such as those enforced by entry exclusion systems (Garcillán-Barcia and de la Cruz, 2008; Julin, 2014) or plasmid-encoded CRISPR-Cas systems (Pinilla-Redondo *et al.*, 2019).

Although the TOL plasmid revealed comparatively lower transfer frequencies within Bregnerød sand filter communities (<10^-3^), it’s transfer within Kerteminde and Herning sand filter communities was relatively high (between 10^-2^ and 10^-1^). Notably, this plasmid is of particular interest in the context of groundwater bioremediation because it naturally carries genes encoding for enzymes involved in the catabolism of toluene and xylenes, which confer its bacterial hosts the ability to degrade several pesticides (Assinder and Williams, 1990). Moreover, RP4 and pKJK5 belong to the IncP group of conjugative plasmids, a diverse family in which some members are also known to harbor pesticide degrading genes (Assinder and Williams, 1990; Dunon *et al.*, 2013; Dealtry *et al.*, 2014).

In contrast to previous work (Li *et al.*, 2018), pKJK5 showed relatively lower transfer frequencies than RP4 (~1 order of magnitude difference) across all water work microbial communities, underscoring the importance of studying plasmid transfer in a case by case manner and for striving to address transfer dynamics in more relevant spatio-temporal settings.

While RP4, pKJK5, and TOL are self-transmissible, RSF1010 does not contain the complete set of necessary genes for conjugation. As such, RSF1010 can only transfer into recipient cells by borrowing components of the conjugative machinery from co-occurring conjugative elements (eg. integrative conjugative elements (ICEs) and conjugative plasmids) through processes known as mobilization and retromobilization (Meyer, 2009; Lee, Thomas and Grossman, 2012). Therefore, monitoring the dissemination of a mobilizable plasmid presents a unique opportunity to measure the intrinsic ability of sand filter communities to mobilize non-self-transmissible plasmids.

Congruent with previous findings, RSF1010 transfer was in the range of 1 order of magnitude lower than RP4 (Klümper *et al.*, 2014), except within the recipient microbial community originating from Herning (Figure 1). The ability of RSF1010 to transfer at high frequencies is remarkable considering its dependency on the availability of compatible conjugation machinery *in trans*. These data indicate a high prevalence of naturally occurring conjugative elements in sand filter communities and demonstrates that plasmid mobilization is an effective gene delivery mode in this environment. Because the conjugative transfer machinery often occupies a large fraction of a plasmid’s genome (Norman, Hansen and Sørensen, 2009), mobilizable plasmids may allow for larger accessory gene cargos, making them particularly suitable vectors for spreading multiple pesticide degrading genes. Moreover, since mobilizable plasmids tend to exist in higher copy numbers (Watve, Dahanukar and Watve, 2010), the increased gene dosage could lead to higher expression levels of bioremediation determinants. Interestingly, Herning sand filter communities showed a higher mobilizing potential than Kerteminde and Bregnerød, suggesting that the presence of compatible conjugative elements may be variable across sand filters. Together, these results highlight the relevance of designing parallel meta-mobilome studies to investigate the indigenous pool of MGEs (Jørgensen *et al.*, 2014; Browne *et al.*, 2020). Such studies would enable a deeper understanding of the factors affecting the plasmid transfer potential of exogenous plasmids within complex microbes.

### Plasmid transfer host ranges across sand filter recipient communities

Although transfer frequency measures provide valuable knowledge regarding the quantitative dissemination potential of plasmids within a given microbial community, they do not inform about the taxonomic permissiveness of communities towards incoming plasmids. In order to explore the transfer host range of the four plasmids across sand filter communities, we isolated the transconjugant populations via cell-sorting (FACS) after mating and characterized them by 16S rRNA gene amplicon sequencing.

Out of all filter mating combinations, a total of 147 distinct transconjugant operational taxonomic units (OTUs) were detected through 16S rRNA sequencing (Figure 2). The transconjugant fractions were largely consisting of taxa from the phylum Proteobacteria, such as several bacteria belonging to the Pseudomonadaceae, Aeromonadaceae and Enterobacteriaceae families (Figure 3a)—consistent with previous reports of the host ranges of the 4 plasmids tested (Mølbak *et al.*, 2003; Klümper *et al.*, 2015; Jacquiod *et al.*, 2017; Li *et al.*, 2018). Interestingly, plasmid transfer was observed into certain Gram-positive taxa, including members of Actinobacteria and Firmicutes (Figure 2), highlighting the exceptional ability of plasmids to cross distant phylogenetic barriers. In agreement with previous studies, trans-Gram transfer appeared to comprise only a small fraction of detected transfer events (Klümper *et al.*, 2015, 2017; Jacquiod *et al.*, 2017; Li *et al.*, 2018) (Figure 2 and 3a, Supplementary Figure S3). Future studies are needed to investigate the extent to which these plasmids can be stably maintained within Gram-positive hosts or whether these bacteria may comprise replicative or transfer dead ends for these MGEs.

**Figure 2.**
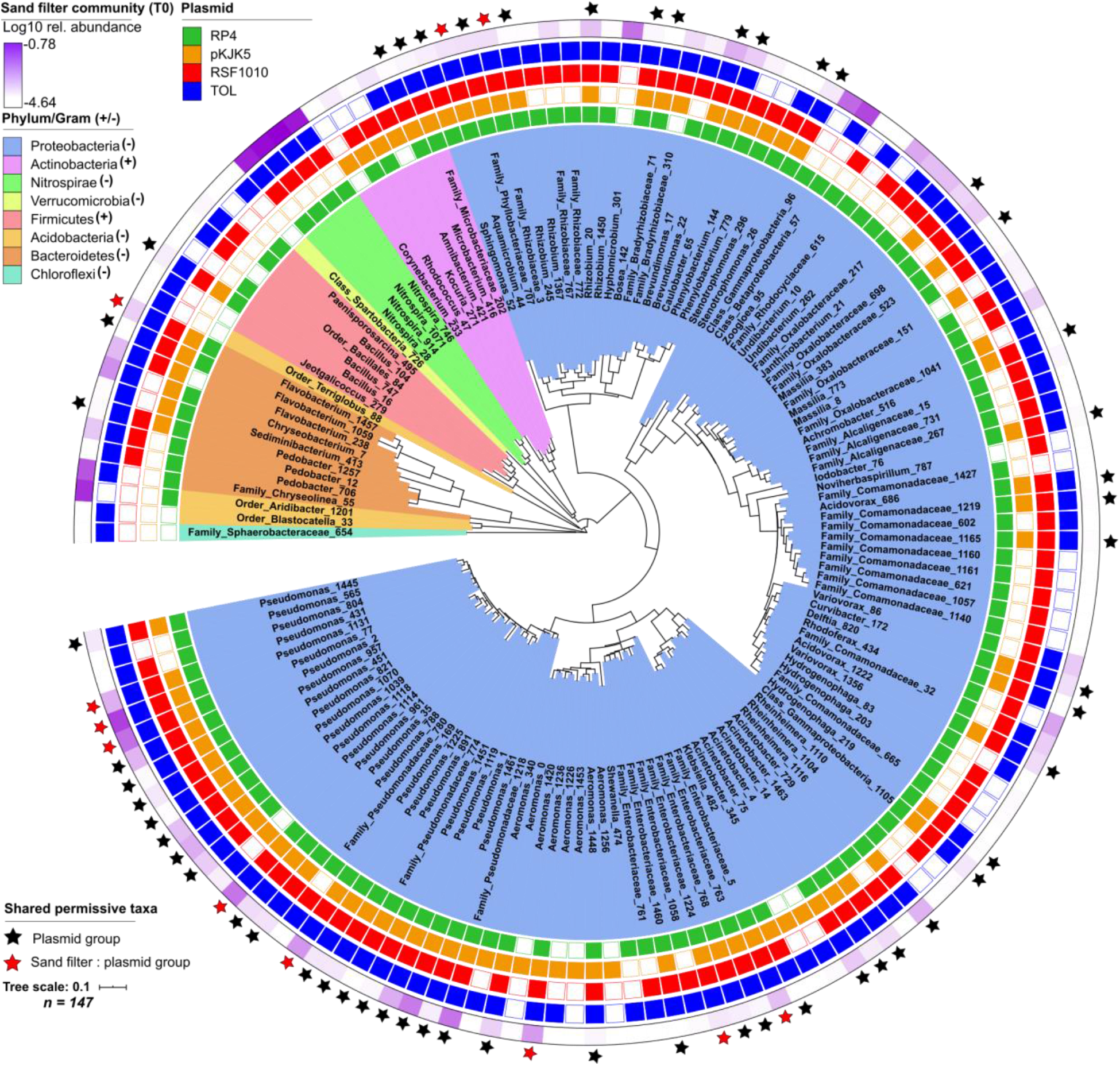
Taxonomic composition of the transconjugant pools across plasmid-donor combinations. Phylogenetic tree showing the identified transconjugant OTUs across all filter matings using *P. putida* as the plasmid donor strain. Only OTUs detected in 2 out of the 3 replicates are displayed. Background colors radiating from the tree indicate the different Phyla to which transconjugants belong, as indicated in the figure legend. For each of the four plasmid-donor combinations (TOL, RP4, pKJK5 and RSF1010) the presence or absence of a given OTU is represented as a colored or empty box, respectively, in the outer concentric lanes. The relative abundances (Log10-transformed) of transconjugant OTUs in the original rapid sand filter communities, herein labeled “T0”, are displayed in the outermost lane (purple). OTUs that were detected in the transconjugant pools across all four plasmid-donor combinations are indicated with a black asterisk. Members which were additionally identified across all three rapid sand filter communities (Herning, Bregnerøed and Kerteminde) are indicated with a red asterisk. The total number of OTUs displayed in the tree is specified in the figure legend.

**Figure 3.**
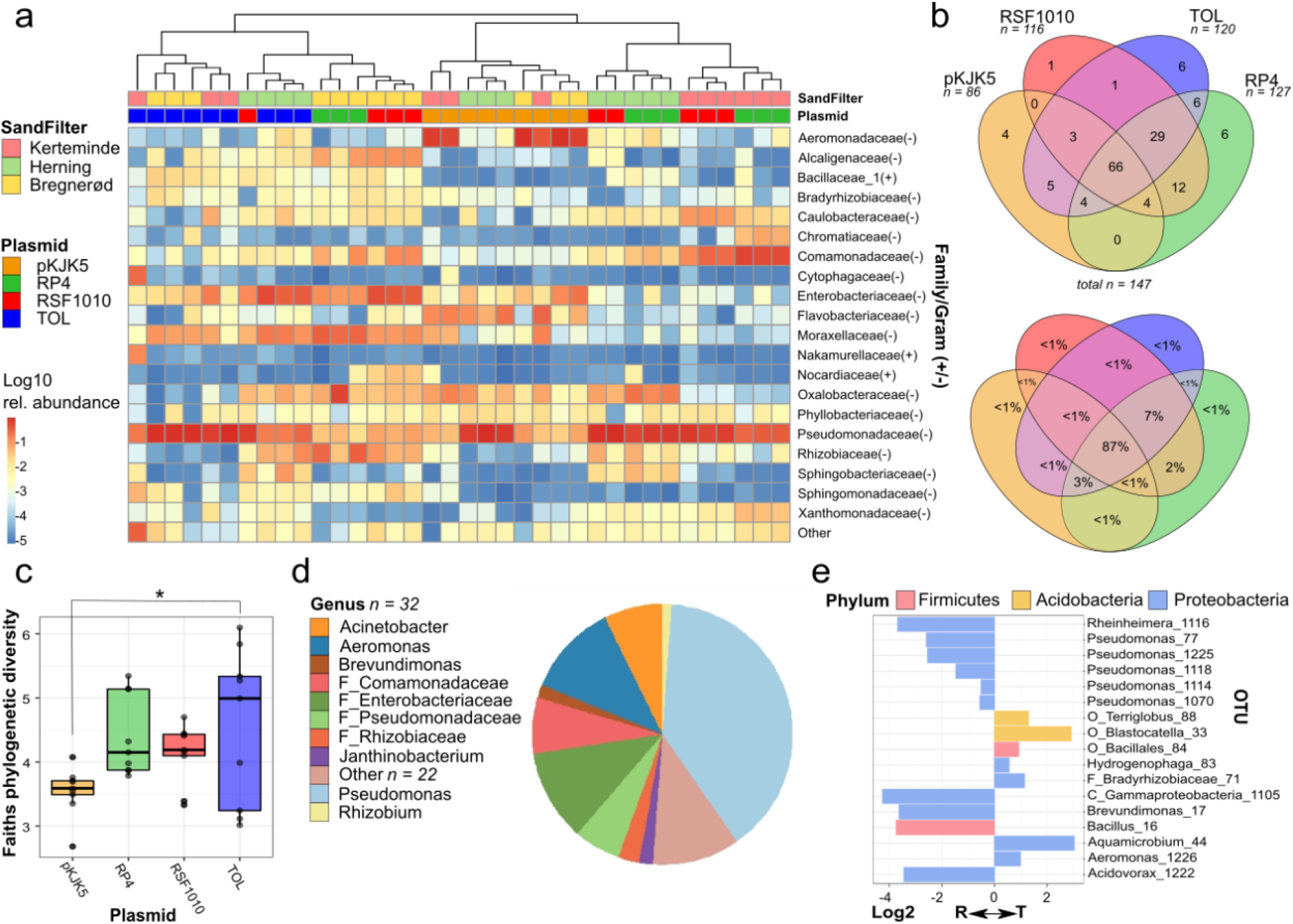
Analysis of the identified transconjugant pools. **a)** Heatmap representing the Log10 relative abundances of the top 20 most prevalent bacterial Families across transconjugant pools. Families found at lower abundance have been grouped under “Other”. The dendrogram shows clustering of samples according to taxonomic abundance, using the Ward method (Murtagh and Legendre, 2014). Filter mating samples are color-coded according to the sand filter community and plasmid used, as indicated in the figure legend (top right). **b)** Venn diagrams displaying the distribution of shared OTUs (top) and their relative abundances (bottom) across all sorted transconjugants populations. The number of OTUs present in each plasmid-specific transconjugant pool are indicated by “n”. **c)** Boxplot showing the Faith’s phylogenetic diversity measure (Faith, 1992) for the 4 plasmid-specific transconjugant pools. The asterisk indicates a significant difference (Tukey honest significant differences, p. adj < 0.05) between TOL and pKJK5. **d)** Pie chart showing the relative abundance distribution of the the 66 OTUs (32 genera) shared across all transconjugant pools (see 3b); 22 genera which were found at relative abundances below 1% are grouped under “Other”. **e)** Barplot showing the Log2 fold change in abundance between the FACS-sorted recipient and transconjugant OTU pools. Only the OTUs which both revealed a significant abundance change (Wilcoxon test with False discovery rate (Benjamini and Hochberg, 1995), p.adj < 0.05) representing 51 out of the total 147 transconjugant OTUs), and a Log2 fold change above 0.5, are displayed. An expanded list can be found in Supplementary Figure S5.

Noteworthy, 66 distinct OTUs pertaining to 32 different genera were detected across all plasmid-donor combinations and comprised 87% of the overall transconjugant pools (Figure 3b). These results reinforce the notion that microbial communities harbor a core super-permissive fraction of bacteria that more readily engage in the uptake (and potential re-transfer) of incoming MGEs (Klümper *et al.*, 2015; Li *et al.*, 2018). In accordance with previous findings (Klümper *et al.*, 2015, 2017; Jacquiod *et al.*, 2017; Li *et al.*, 2018; Cyriaque *et al.*, 2020), the core permissive taxa mainly consisted of different Proteobacteria, including genera such *Pseudomonas*, *Acinetobacter*, *Aeromonas*, *etc.* and members of the *Enterobacteriaceae* and *Rhizobiaceae* families (Figure 2, Figure 3d, and Supplementary Figure S3). These results suggest common strategies in the promiscuity of certain taxa towards foreign incoming DNA.

The Faith’s phylogenetic diversity index was evaluated across plasmid-specific transconjugant pools as a proxy for taxonomic breadth of transfer (Figure 3c). Overall, our analysis revealed similar transfer ranges for the different plasmids at the conditions tested, except for pKJK5, which appeared to show a comparatively lower transconjugant OTU richness (Figure 3b and 3c)—significantly lower than that of the TOL plasmid (Tukey honest significant differences, p. adj < 0.05). Interestingly, the plasmid RSF1010 displayed a relatively broad transfer host range (Figure 3c), transferring into 116 distinct OTUs despite being non-self transmissible (Figure 3b). Our results thus support the idea that because mobilizable plasmids can often be shuttled by diverse type IV secretion system machineries, they can disseminate remarkably across microbial communities. Notably, these exceptional properties have been proposed to allow mobilizable plasmids access to even broader taxonomic host ranges than self-transmissible plasmids under natural conditions (Meyer, 2009; Klümper *et al.*, 2014).

Importantly, certain taxa that were represented in the filter-mated recipient communities at low abundances were found to be relatively enriched in the corresponding transconjugant pools, indicating their high permissiveness towards incoming plasmids. These include members of *Aquamicrobium*, *Blastocella*, and *Terriglobus*, among others. On the other hand, certain abundant recipient genera, such as *Acidovorax*, *Reheinheimera*, *Brevindimonas*, *Bacillus*, and certain *Pseudomonas* were poorly represented in the transconjugant pools (Figure 3e, Supplementary Figure S5).

Altogether, these results emphasize the intricate dynamics surrounding plasmid-host interactions and the need for a deeper characterization of the recipient and transconjugant pools. Given that both innate and adaptive barriers against foreign invading genetic elements are extremely diverse and heterogeneously spread across bacterial taxa (Thomas and Nielsen, 2005), differences in plasmid transfer between communities and community members (even among closely related taxa) are expected. Future studies will benefit from the advent of complementary procedures to 16S rRNA gene amplicon sequencing, such as advances in sequencing single-cell amplified genomes (SAGs) (Tolonen and Xavier, 2017). Indeed, comparative genomic analyses of recipient and transconjugant cell SAG data could provide insights into the genetic factors influencing the promiscuous or refractory nature of bacteria towards incoming plasmids.

A clear distance between the indigenous sand filter communities (T0) and the recipient communities from the filter matings was observed (Supplementary Figure S4, PERMANOVA on weighted unifrac distance, R2 = 0.71, p = 0.001). This is likely due to a large fraction of the total indigenous bacteria from the sand filters not being able to grow under the conditions tested Additionally, since it is expected that certain transconjugant cells may miss detection due to plasmid loss or inadequate GFP expression levels, it is likely that the natural transfer host ranges of these plasmids are even broader than reported here. These considerations serve as a reminder that the paradigms derived from our model system may not faithfully extend to the natural environment, where the microbial taxonomic diversity and physicochemical conditions are significantly different. However, given that frequent and broad-range transfer detected within the cultivable fraction of all the sand filter communities investigated here (Figure 1 and 2, Supplementary Figure S6), we conclude that rapid sand filters likely constitute environments with high permissibility potentials towards incoming plasmids.

Furthermore, our findings are relevant in the assessment of the risks posed by the spread of plasmid-borne antimicrobial resistance (AMR) determinants within potable water systems. The presence of AMR genes on environmental plasmids is widely acknowledged and since aquatic environments represent natural drainage spots for environmental microbes, aquifers may constitute unrecognized reservoirs for MGEs encoding AMR. Indeed, AMR genes have already been reported to be commonly present in groundwater ecosystems (O’Dwyer *et al.*, 2017; Harb *et al.*, 2019). While plasmid transfer dynamics within aquatic environments may be hampered by reduced cell-to-cell contact events, surface-associated sand filter microbial communities may constitute dissemination hotspots for AMR-encoding plasmids. Standard water quality assessments are centered around the strict absence of coliforms (World Health Organization, 2011), yet the presence of innocuous bacteria carrying antibiotic resistance plasmids is currently overlooked. Our data thus highlights the necessity for future investigations of the risks associated with the presence and transfer of AMR plasmids within potable water treatment facilities, as well as an evaluation of potential measures required to mitigate this phenomenon.

## Conclusions

The contamination of groundwater ecosystems with anthropogenic pollutants (e.g. pesticides) has severe environmental repercussions that challenge the production of potable water. Notably, groundwater-fed water works are not equipped with the natural pesticide-degrading capabilities required to face this growing concern. Cell-based bioaugmentation practices have been proposed to mitigate this problem, yet these efforts are typically limited due to out-competition of the inoculated strains by indigenous microbes (ecological barrier effect). On the other hand, plasmid-based bioaugmentation approaches have the potential to enhance the degradative competence of the already established, ecologically competitive, autochthonous microbial communities. This study revealed the significant ability of natural plasmids to transfer at high frequencies and across distantly related taxa within groundwater-fed rapid sand filter communities in the absence of plasmid selection, indicating their suitability as vectors for the spread of pesticide degrading determinants in water purification plants. Furthermore, our data shows that mobilizable plasmids, despite being non-self transmissible, can disseminate widely—comparable to the dissemination potentials attained by certain broad-host range conjugative plasmids. Future work is required to assess the biotechnological applicability and long-term maintenance of exogenous plasmids within sand filter communities.

## Supplementary Information

**Supplementary Fig. S1.**
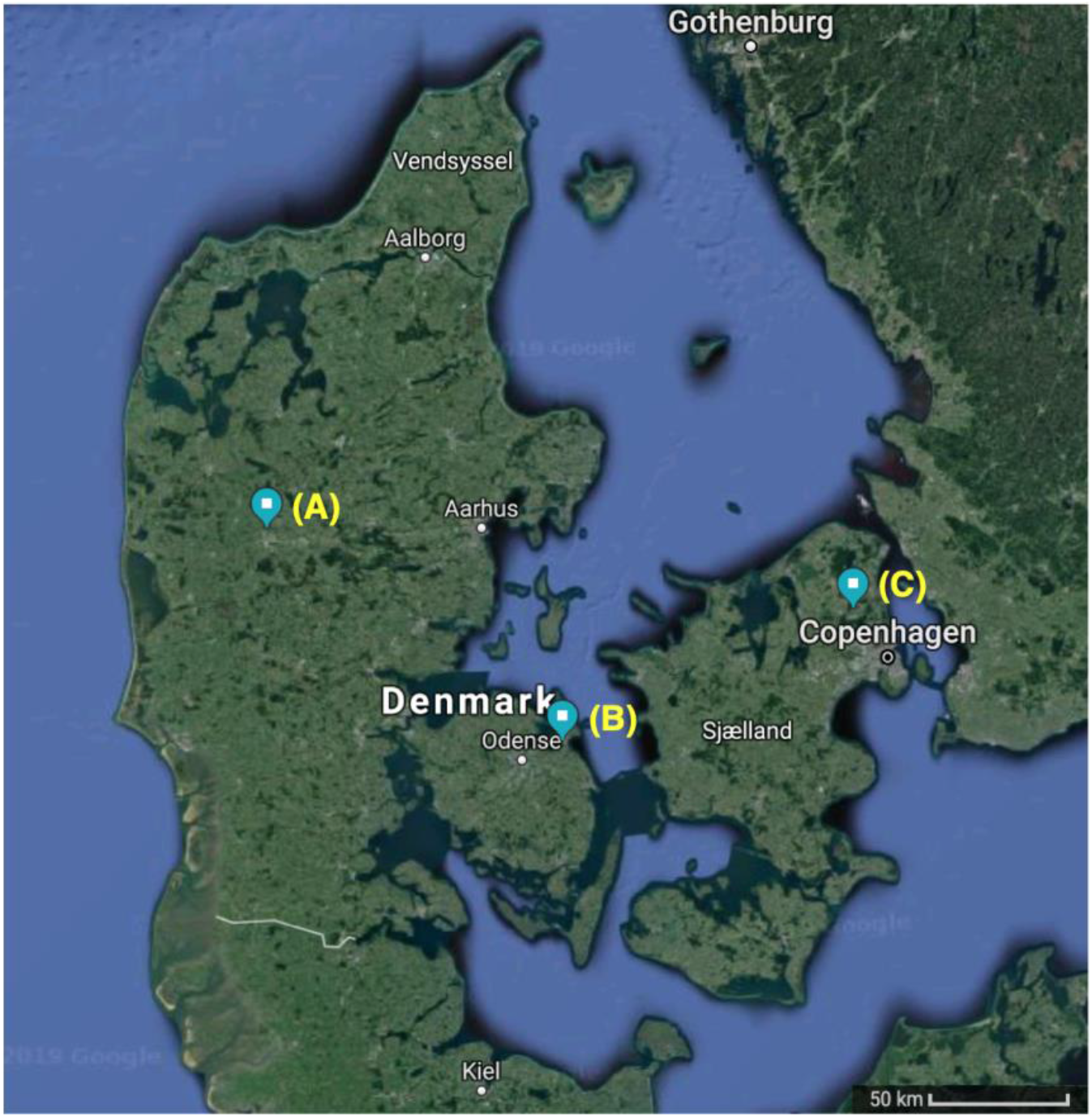
sand filter sampling locations. Google map caption indicating the locations of the waterworks in Denmark from which sand filter samples were taken: (A) Herning; (B) Kerteminde; (C) Bregnerød.

**Supplementary Fig. S2.**
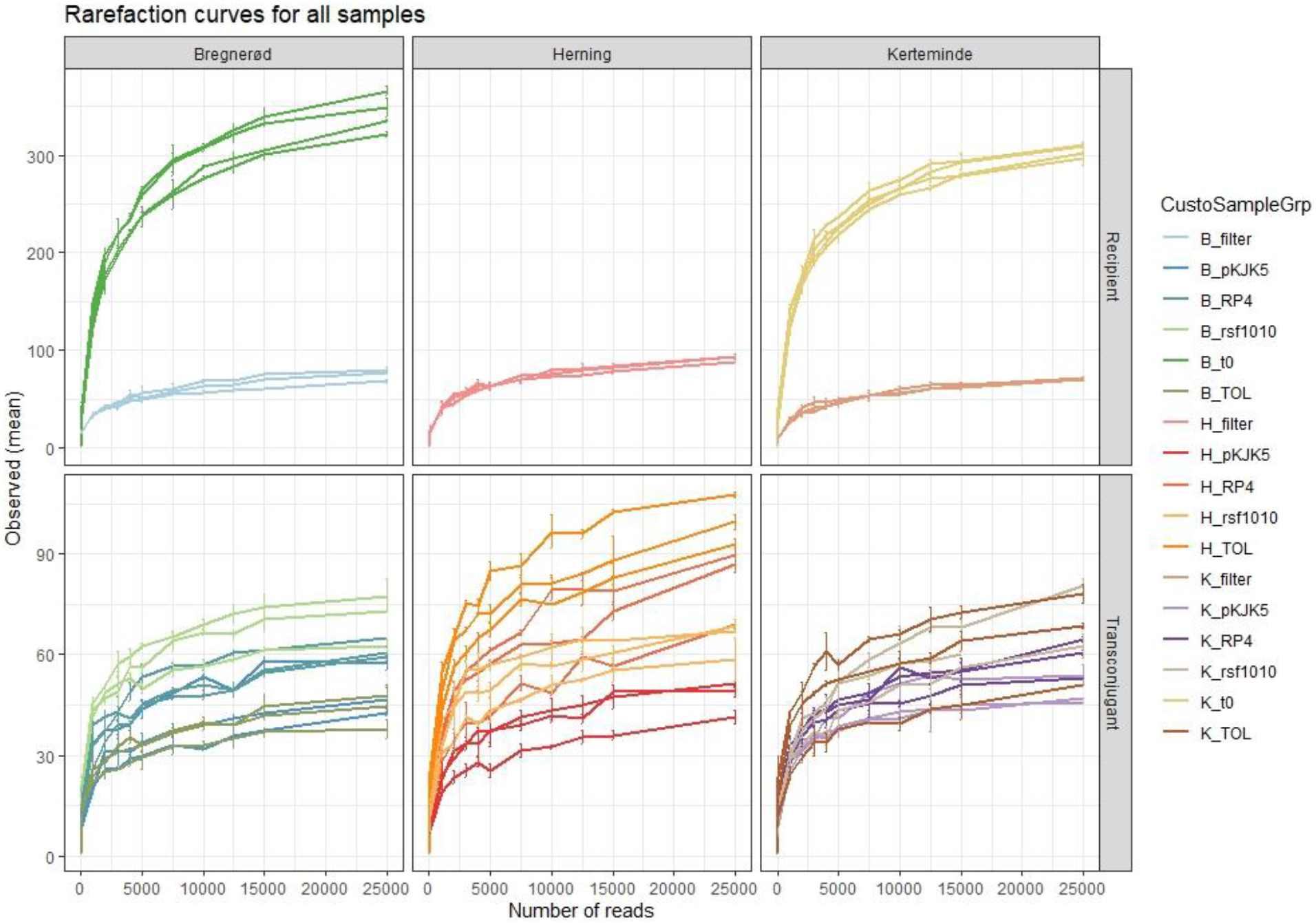
Rarefaction curves of the 16S rRNA gene sequencing profiles for each of the studied samples. Curves show the observed OTUs at 97% similarity in each sample as a function of the number of 16S rRNA gene sequencing reads (X-axis). Colors indicate the sample groups indicated in the figure legend; recipient communities (original; “t0”, and sorted post-mating; “filter”) and transconjugant (sorted post-mating; “filter”) samples that are separated by plasmid (TOL, RSF1010, pKJK5, and RP4) and sand filter location: Bregnerød (B), Kerteminde (K), and Herning (H).

**Supplementary Figure S3.**
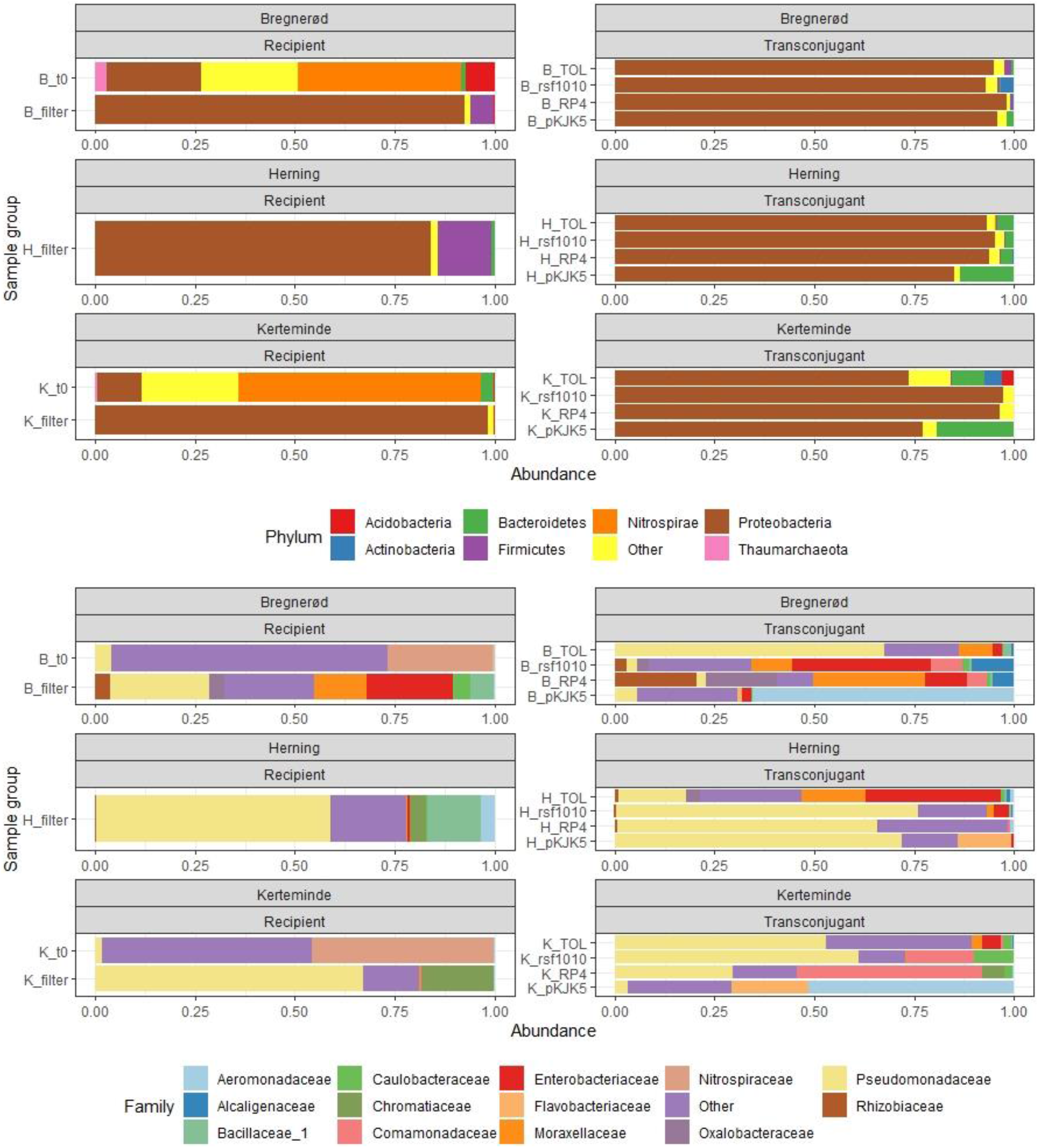
Phylogenetic composition of the samples analysed in this study. Bar plots show the relative abundances of taxa at the Phylum (top) and Family (Bottom) levels for all sequenced samples: original sand filter communities (“t0”, average of 4 replicates) and FACS-sorted recipients (“Filter”, average for 3 replicates) and transconjugant pools (“plasmid”, average of 3 replicates), grouped by sand filter water work location and plasmid-donor combination. Taxa below 0.1% relative abundance, for phylum level, and 1% for family level, have been grouped into “other”. Raw data found in Supplementary Table S2.

**Supplementary Figure S4.**
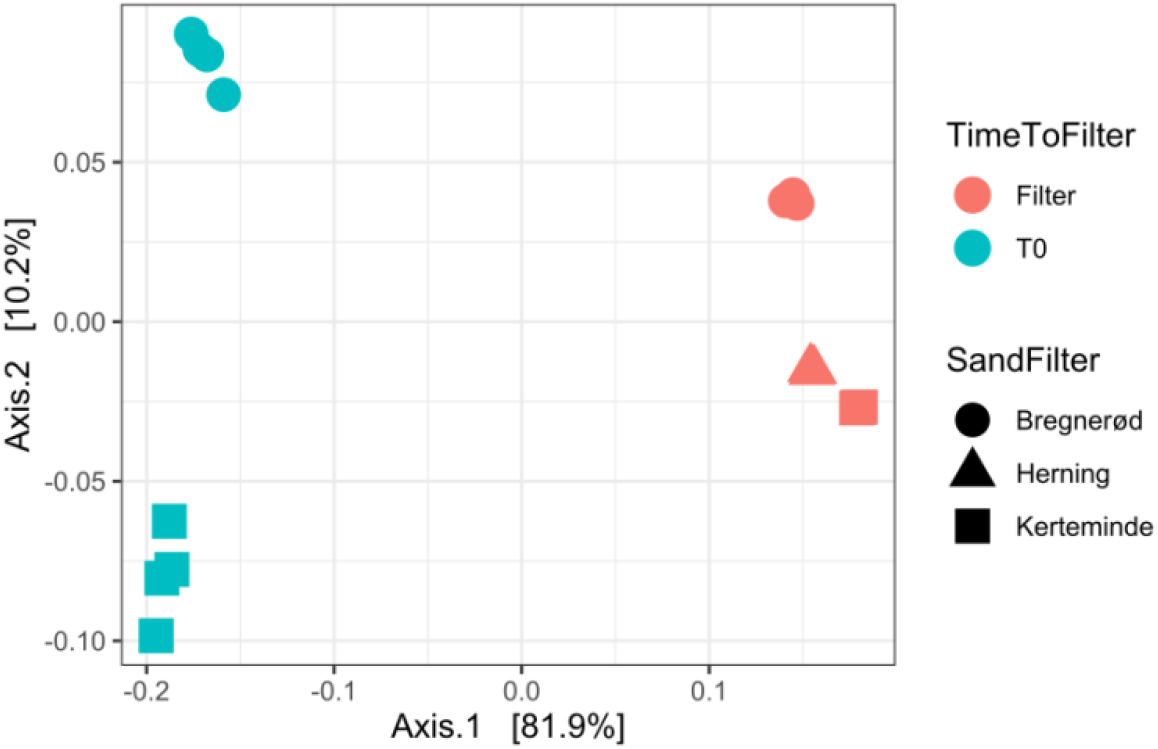
Culturing rapid sand filter bacteria substantially affects the original microbial community composition. PCoA of the weighted unifrac distance between the recipient sand filter communities, as indicated by icon shape, and recipient community: original sand community; “T0”, and sorted post-filter mating; “Filter”, shown in blue and red color, respectively.

**Supplementary Figure S5.**
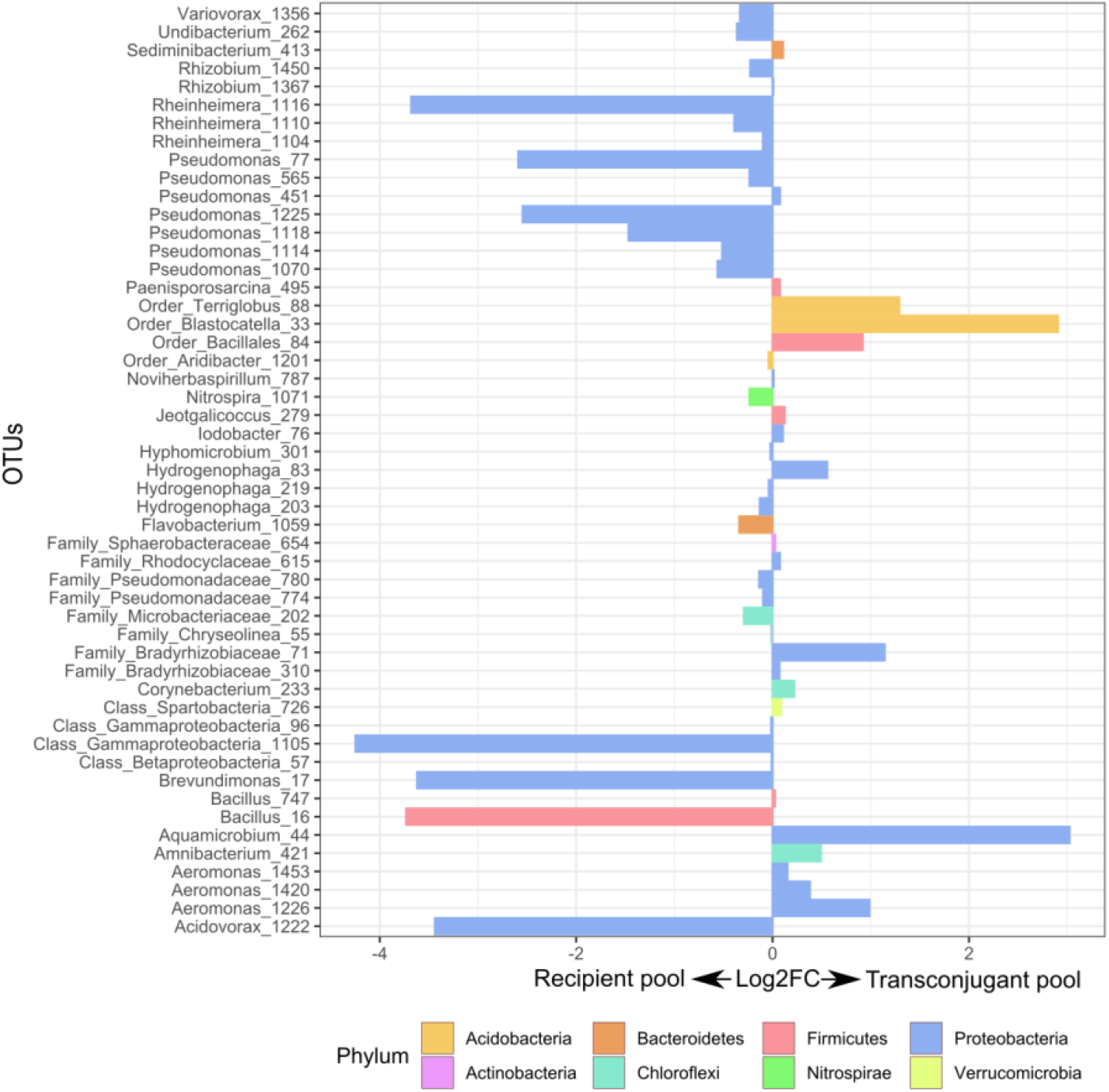
Log2 fold abundance change between the FACS-sorted recipient and transconjugant OTU pools. Only OTUs which displayed a significant abundance change using the Wilcoxon test; adjusted for multiple testing (p.adj < 0.05) are displayed; 51 out of the total 147 transconjugant OTUs.

**Supplementary Figure S6:**
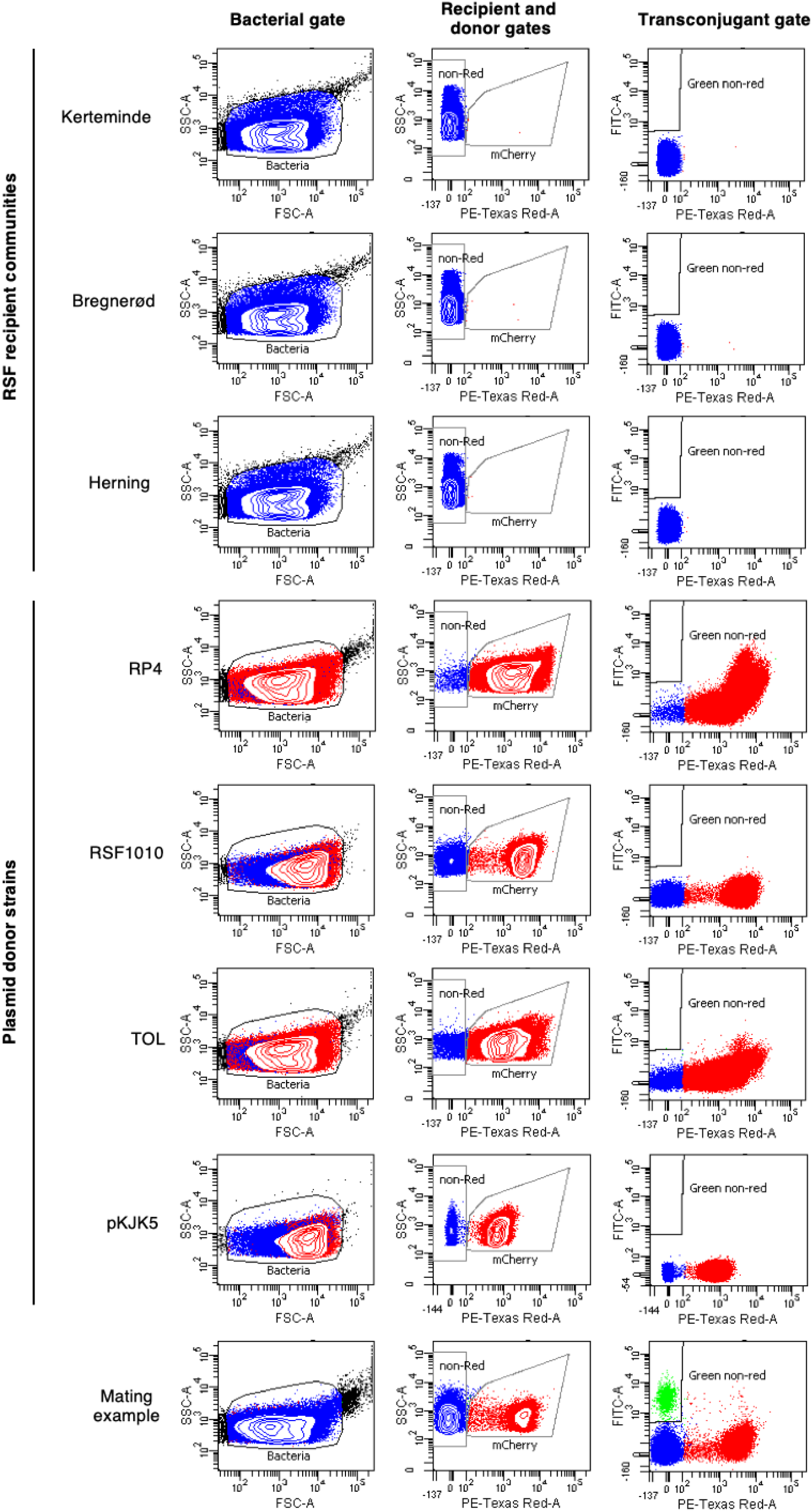
Overview of the flow cytometry and FACS gates employed for counting and sorting plasmid-donor, recipient and transconjugant cells. The different sand filter communities and plasmid donor strains that were used in this work were grown on solid-surface filters alone and served as negative controls for transfer during filter matings. The resultant Log(10)-scaled scatter plots were employed to design an appropriate gating strategy; (i) bacterial cells (left column, FSC-A vs. SSC-A; in blue); (ii) red fluorescent donor cells and non-red recipients (middle column; FSC-A vs. PE-Texas-Red-A, in red and blue, respectively); (iii) green non-red fluorescent transconjugant cells (right column, FITC-A vs. PE-Texas-Red-A). Bregnerød recipient community challenged with *P. putida* carrying RP4, replicate 2, is displayed in the last row as an example of a filter mating outcome and to show the suitability of the designed transconjugant cell gate.

**Supplementary Table S1. Flow cytometry cell counts and transfer efficiency calculations. (Separate Datasheet)**

**Supplementary Table S2: 16S rRNA gene sequencing data generated in this study.**

**(Separate Datasheet)**

## Acknowledgements

Financial support for this project was provided by the Innovation Fund Denmark, Trojan Horse Project [#5157-00005B], the Independent Research Fund Denmark, InTrans Project [#8022-00322B] and Joint Programming Initiative-Antimicrobial Resistance (JPI-AMR), DARWIN Project, [#7044-00004B]. J.R. was supported by the Novo Nordisk Foundation [#NNF17OC0025014].

## Author contributions

S.M., S.J.S, and R.P-R. conceived and planned the experiments. S.M. and L.D.C carried out the filter mating experiments. R.P-R. performed FACS analyses and sequencing. A.K.O and J.N. analysed the sequencing data with the help of J.R. R.P-R. wrote the article with input from all authors.

